# RK-33, a small molecule inhibitor of host RNA helicase DDX3, suppresses multiple variants of SARS-CoV-2

**DOI:** 10.1101/2022.02.28.482334

**Authors:** Farhad Vesuna, Ivan Akhrymuk, Amy Smith, Paul T. Winnard, Shih-Chao Lin, Robert Scharpf, Kylene Kehn-Hall, Venu Raman

## Abstract

SARS-CoV-2, the virus behind the deadly COVID-19 pandemic, continues to spread globally even as vaccine strategies are proving effective in preventing hospitalizations and deaths. However, evolving variants of the virus appear to be more transmissive and vaccine efficacy towards them is waning. As a result, SARS-CoV-2 will continue to have a deadly impact on public health into the foreseeable future. One strategy to bypass the continuing problem of newer variants is to target host proteins required for viral replication. We have used this host-targeted antiviral (HTA) strategy that targets DDX3, a host DEAD-box RNA helicase that is usurped by SARS-CoV-2 for virus production. We demonstrated that targeting DDX3 with RK-33, a small molecule inhibitor, reduced the viral load in four isolates of SARS-CoV-2 (Lineage A, and Lineage B Alpha, Beta, and Delta variants) by one to three log orders in Calu-3 cells. Furthermore, proteomics and RNA-seq analyses indicated that most SARS-CoV-2 genes were downregulated by RK-33 treatment. Also, we show that the use of RK-33 decreases TMPRSS2 expression, which may be due to DDX3s ability to unwind G-quadraplex structures present in the TMPRSS2 promoter. The data presented supports the use of RK-33 as an HTA strategy to control SARS-CoV-2 infection, irrespective of its mutational status, in humans.

## Introduction

The causative agent of the worldwide COVID-19 pandemic is the Severe Acute Respiratory Syndrome Coronavirus 2 (SARS-CoV-2). Vaccines are generally considered as the first line of defense against such a large-scale viral spread. Basically, vaccines prime the host’s immune system, which then coordinates a direct attack against specific virions. Indeed, rapid deployment of several safe COVID-19 vaccines has proven very effective in attenuating infections and limiting deaths. Nevertheless, as with all viruses, SARS-CoV-2 has been evolving/mutating and variants that are more transmissible have emerged ^1, 2^. Moreover, it is possible that the continuous evolution of SARS-CoV-2 will result in a vaccine resistant variant(s) ^2^ even as the efficacy of vaccines is waning ^3–5^. Accordingly, SARS-CoV-2 will continue to have a major impact on the future of public health if development of alternative therapeutic strategies is not pursued.

Direct-acting antiviral agents (DAAs) and those that are host-targeted antivirals (HTAs) have been developed into effective alternatives to vaccine resistance or for use in combination with vaccinations ^6^. Here, we have used a HTA directed against the human host DEAD (Asp, Glu, Ala, Asp)-box RNA helicase, DDX3 (DDX3X) and demonstrated a potential of providing an anti-SARS-CoV-2 therapy. We developed a small molecule inhibitor of DDX3, RK-33, that is designed to be a competitive inhibitor of DDX3’s ATP binding site and has been demonstrated to abrogate its RNA helicase function ^7–9^ which is required for the translation of complex structured 5`’`capped mRNAs. Our rationale was based on evolutionary studies of SARS-CoV-2, which have revealed that it is closely related to SARS-CoV with a similar sized genome that has nearly identical organization and hence, replication ^10–15^. SARS-CoV-2 replication initially proceeds through a 5`’`cap-dependent translation using the host’s translation complexes ^16^. Consequently, SARS-CoV-2 also has a 5`’`capped mRNA genome that requires cap-dependent translation ^17^. Given the strong evidence that DDX3 participates in 5`’`capped translation where it forms functional complexes with several eukaryotic translation initiation co-factors including eIF4E, eIF4F, eIF4G and eIF3 ^18–21^ and is a required functioning unit in promoting translational as a part of the 80S translation initiation complex ^18^, it follows that DDX3 would be utilized in the translation of the SARS-CoV-2 genome. Consistent with this line of reasoning, recent proteomics analyses found DDX3 colocalized with SARS-CoV-2 viral particles and the latter required DDX3 for virion production ^22^, DDX3 interacted with SARS-CoV-2 RNA ^23^, and that DDX3 was part of the SARS-CoV-2 interactome ^24^.

Similarly, DDX3 also participates in the resolution of both RNA and DNA G-quadraplex structures and such structures have been predicted to form in the SARS-CoV-2 genome ^25–27^, in the promoter region of the host TMPRSS2 gene ^28^, as well as in the 5`’`UTR of the TMPRSS2 transcript. TMPRSS2 expression is also required for SARS-CoV-2 entry into cells ^29–31^. In addition, three related lines of evidence indicate that DDX3 can be considered as a prime host protein to target ^32^: a) several viruses (18 species from 12 genera and 11 families) have been demonstrated as requiring DDX3 for efficient virion production; b) investigators have actively pursued the targeting of DDX3 with small molecule inhibitors as an antiviral strategy ^33–42^; and c) RK-33 has effectively been used to target DDX3 and abrogate virion production in Dengue, West Nile, Zika, Respiratory Syncytial, and human Parainfluenza Type-3 viral infections ^40^.

Our results indicate that regardless of the SARS-CoV-2 isolate (Lineage A, and Lineage B Alpha, Beta, and Delta variants), RK-33 treatment of Calu-3 cells infected with these isolates display significantly reduced viral load by one to three log orders. Consistent with this finding, proteomics and RNA-seq analyses of the infected/RK-33 treated cells demonstrated the downregulation of most of the SARS-CoV-2 proteins and genes and host TMPRSS2 expression was decreased as well. Collectively, the data obtained indicates that the use of RK-33 to abrogate host DDX3 functions is a viable option for the treatment of SARS-CoV-2 infection as well as subsequent variants that may arise.

## Results

### RNA-seq demonstrates RK-33 treatment downregulates multiple viral transcripts and alters the host transcriptional landscape

To study the effect of RK-33 and SARS-CoV-2 on the human transcriptome, we performed RNA-seq on isolated RNA from uninfected Calu-3 cells treated with DMSO (DMSO control), cells treated with RK-33 (RK-33 treated DMSO control), cells infected with SARS-CoV-2 and treated with DMSO (Virus), and cells infected with SARS-CoV-2 and treated with RK-33 (RK-33 treated virus). A non-toxic dose of RK-33 (5 μM) was used for these studies and cells were infected at a MOI of 0.1. All samples were collected at 48 h post infection (hpi).

RNA-Seq was mapped separately to human genome and SARS-CoV-2 genome. Principal Components Analysis (PCA) of host RNA showed a good separation between the infected samples and the control treated samples, while the RK-33 treated samples were clustered together. As expected, RK-33 and SARS-CoV-2 infection were the drivers for sample segregation (Figure 1a and S1). Hierarchical clustering separated all RNA samples according to treatment with the viral infected cells being the most different compared to the other three groups (Figure 1b). RNA of most SARS-CoV-2 genes (ORF1ab, S, ORF3a, E, M, ORF7a, ORF7b, ORF8, N, ORF10) demonstrated a reduction when treated with RK-33 compared to virus infected control cells (Figure 1c). Volcano plots were produced to compare host RNA expression between viral infected RK-33 treated vs untreated cells (Figure 1d) and viral infected vs uninfected cells (Figure 1e). Many of the genes that were upregulated by the virus were significantly downregulated by RK-33 treatment. We studied genes that were significantly dysregulated (>1.1 FC, p<0.05) by RK-33 treatment of virally infected Calu-3 cells (Infected RK-33 vs Infected DMSO), by virus infection of the cells (Infected DMSO vs Uninfected DMSO), and with the control (Uninfected RK-33 vs Uninfected DMSO). The gene sets were plotted on a Venn diagram (Figure 1f), and show the largest overlap exists between the RK-33 treated virally infected cells and the virally infected cells, indicating the widespread effects of SARS-CoV-2 on the genetic landscape. Reactome pathway analysis ^43, 44^ revealed that the top five pathways enriched in viral infected DMSO treated control cells were Metabolism of proteins, Gene expression (transcription), Metabolism, RNA Pol II transcription, and Translation (Figure 1 g). The top five Reactome pathways enriched in viral infected RK-33 treated cells were Metabolism of proteins, Metabolism, Post-translational protein modification, Gene expression (transcription), and RNA Pol II transcription (Figure 1 h). It is important to note that host translation is less significantly impacted after cells are treated with RK-33. Gene set enrichment analysis of the uninfected cells and RK-33 treated virus infected cells are shown in Figures S2 and S3. Analysis of genes dysregulated by SARS-CoV-2 infection and RK-33 treatment expectedly showed a larger effect (Figure S3). We also compared our gene set with published data ^45^ which showed a large commonality both in Calu-3 gene sets and in the deceased COVID-19 lung tissue set (Figure S4). In addition, RNA-seq QC results indicated that all samples had around 45 million clean reads, but alignment QC indicated SARS-CoV-2 infected Calu-3 cells had around 33% unique matches in the human genome compared to the rest of the samples at 88% (Figure S5). This is an important result, as it is known that the Nsp1 degrades host RNA so that it can hijack the host cell ^46^ . This reversion of viral to host RNA in the RK-33 treated SARS-CoV-2 infected cells demonstrates the role of DDX3 in SARS-CoV-2 replication and the importance of targeting DDX3 by RK-33.

**Figure 1.**
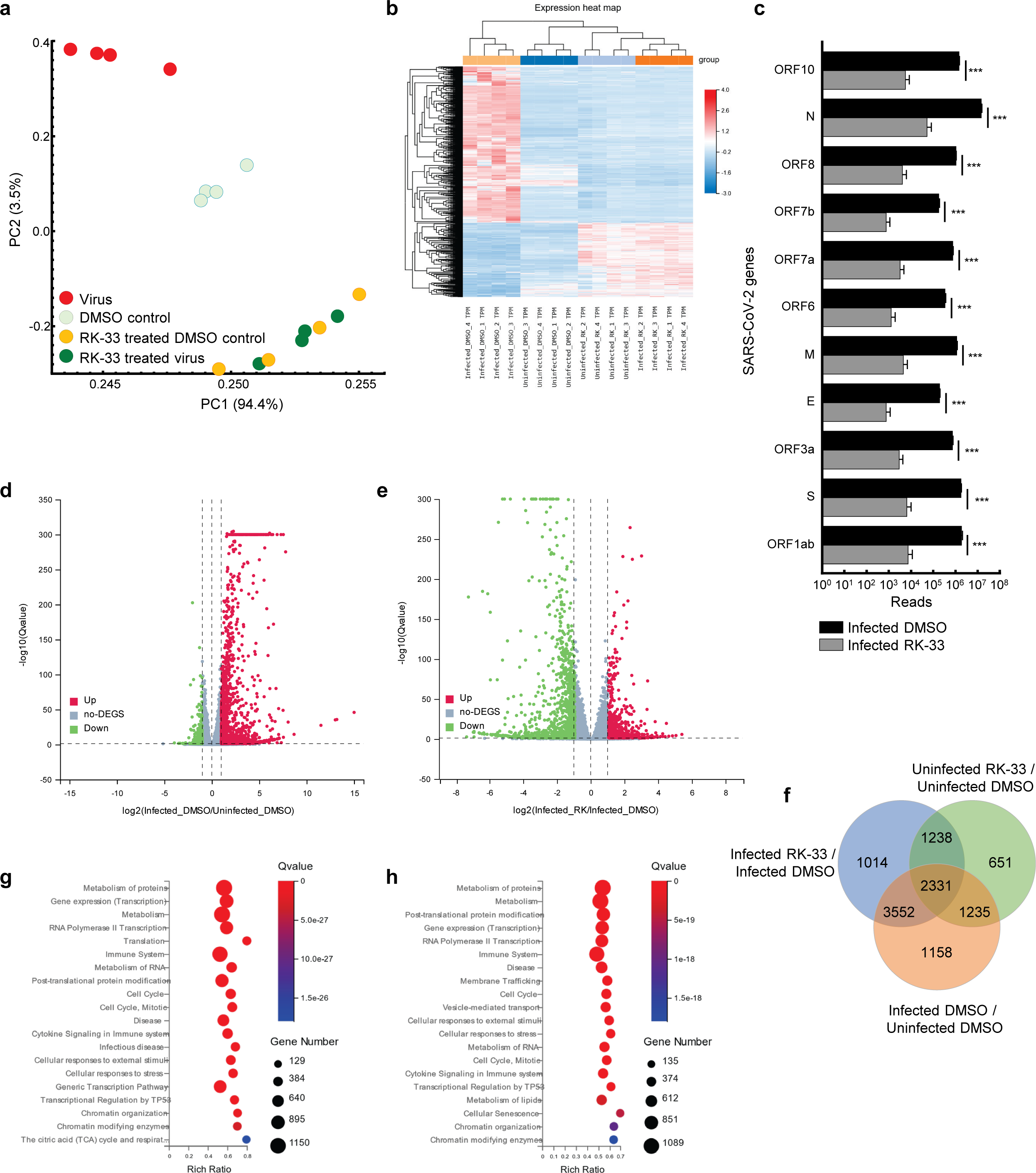
**Transcriptomics of viral infected Calu-3 cells** a. PCA plot of RNA-seq data. PC1 accounts for 94% of the variation between the sample clusters. b. Dendrogram and Heat map of RNA-seq data showing relationship between the various samples. Dendrogram was constructed using z-scores of TPM values of all RNA-seq sample reads. Horizontal sorting was by cluster order. c. Histogram of SARS-CoV-2 RNA that are downregulated by RK-33 treatment of virally infected Calu-3 cells. d. Volcano plot displaying effect of virus infection on Calu-3 cells. Genes differentially regulated in virally infected untreated samples were compared to uninfected samples. The X-axis represents the log2 fold change of the difference, and the Y-axis represents the -log10 significance value. Red represents DEG upregulated (>2-fold change, green represents DEG downregulated, and gray represents non-DEGs. e. Volcano plot displaying effect of RK-33 on virus infected Calu-3 cells. f. Bubble plot displaying significantly enriched (Q<0.05) Reactome pathways affected by SARS-CoV-2 infection of Calu-3 cells. X-axis displays the enrichment ratio which indicates more enrichment towards the right. Larger bubbles indicate more genes. g. Reactome enrichment pathways affected by RK-33 treatment of virally infected Calu-3 cells displayed as a bubble plot.

### Proteomics analysis indicated RK-33 treatment suppresses multiple viral proteins and alters the host response to infection

To study changes in the human and viral proteome after infection with SARS-CoV-2 and RK-33 treatment, we performed proteomics analysis on SARS-CoV-2 infected Calu-3 cells treated with DMSO solvent control (Virus) and on the RK-33 treated and virus infected Calu-3 cells (RK-33 treated virus) as described above. Other controls used were uninfected Calu-3 cells treated with RK-33 (RK-33 treated DMSO control) and Calu-3 cells treated with DMSO (DMSO control). PCA showed clear separation between virally infected and uninfected cells as well as virally infected cells treated with DMSO, and virally infected cells treated with RK-33 (Figure 2a and S6). This indicated that RK-33 and viral infection were the major factors driving the segregation of samples on the PCA plot. Next, we determined the clustering pattern of the untreated and RK-33 treated host cells as well following virus infection. As shown in the dendrogram and heat map (Figure 2b and S7), the virus infected Calu-3 samples and the uninfected samples clustered separately while the RK-33 treated virus infected samples as well as uninfected RK-33 samples clustered amongst each other.

**Figure 2.**
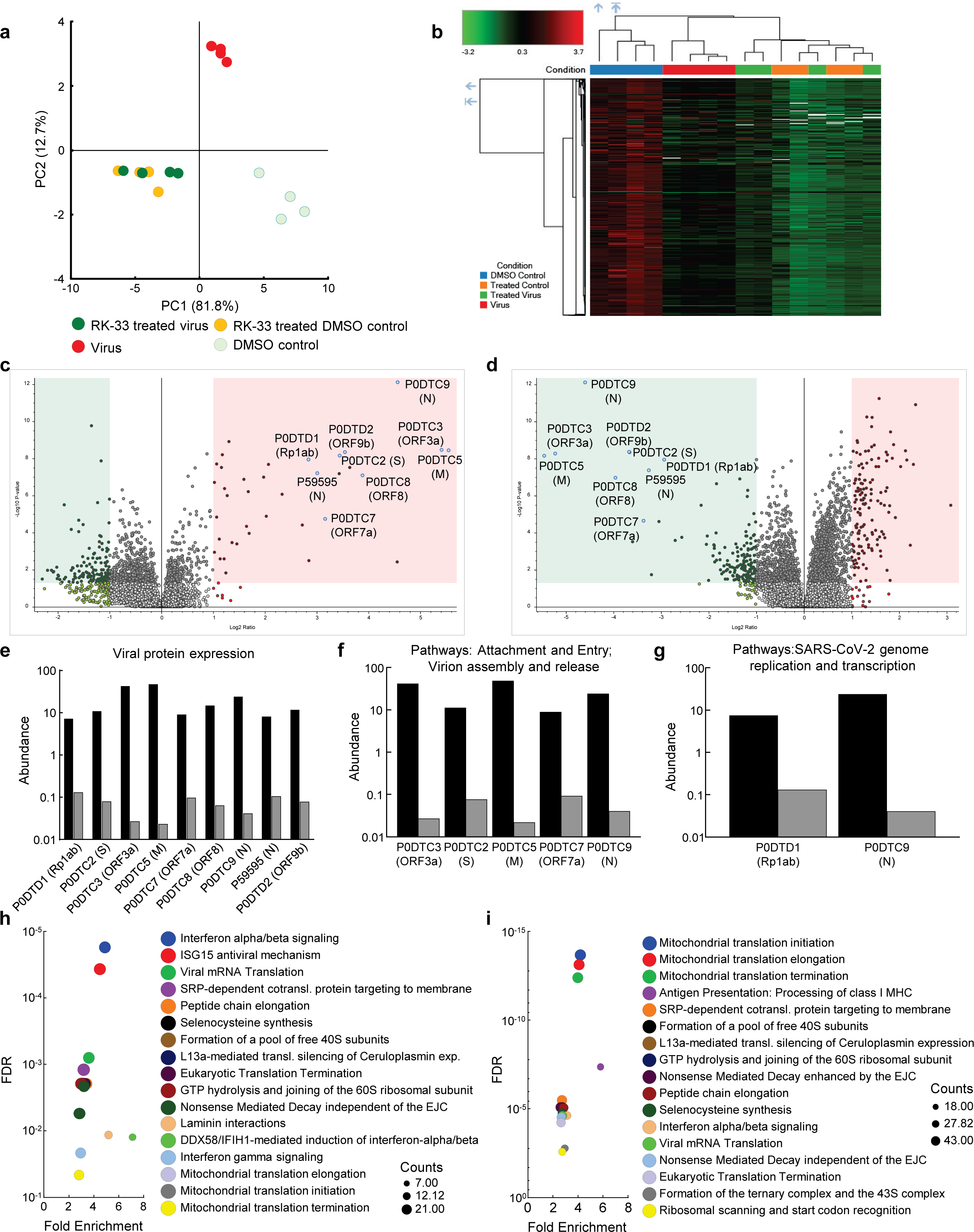
**Proteomics of host and viral proteins from Calu-3 viral infected cells** a. Protein abundances from all samples were analyzed and plotted on a PCA plot. PC1 contributes to 82% and PC2 contributes to 13% of the variation. b. Dendrogram and heat map of proteins as estimated by Euclidean clustering and median linkage. Samples were scaled after clustering. c. Volcano plot displaying SARS-CoV-2 proteins that were differentially regulated following virus infection of Calu-3 cells. Proteins (genes) are labeled. Red shading includes proteins that were significantly upregulated (p<0.05, and 1.1-fold change). Green shading indicates proteins that were significantly downregulated (p<0.05, and -1.1-fold change). d. Volcano plot displaying effect of RK-33 on SARS-CoV-2 proteins in virally infected Calu-3 cells. e. Histogram displaying quantification of viral protein abundances showing effect of RK-33 on SARS-CoV-2 proteins. f. Histogram indicating proteins from Reactome pathways attachment and entry, and virion entry and release affected viral proteins that are downregulated by RK-33 treatment of virally infected cells. g. Histogram of proteins from pathway SARS-CoV-2 genome replication and transcription that are downregulated by RK-33 treatment. h. Bubble chart displaying significant (FDR<0.05) Reactome enrichment of pathways in SARS-CoV-2 infected Calu-3 cells. i. Reactome enrichment pathways of RK-33 treated virally infected cells displayed as a bubble plot.

To study how RK-33 treatment affected Calu-3 infected cells, we plotted the data using Volcano plots (Figures 2c-d). As expected, most of the SARS-CoV-2 proteins (P0DTC2 spike glycoprotein, P0DTC3 ORF3a, P0DTC5 membrane protein, P0DTC8 ORF8, P0DTC9 nucleoprotein, P0DTD1 replicase polyprotein 1ab, P0DTD2 ORF9b, P59595 nucleoprotein) were significantly upregulated (>7.5 FC) after infection of Calu-3 cells. RK-33 treatment significantly downregulated (>8 FC) all these viral proteins in Calu-3 cells. Changes in protein abundance are displayed in Figure 2e-g.

Subsequently, we performed pathway analysis using Reactome to compare RK-33 treated infected cells with control infected cells. The five most significant pathways that were enriched following SARS-CoV-2 infection were interferon alpha/beta signaling, ISG15 antiviral mechanism, viral mRNA translation, SRP-dependent cotranslational protein targeting to membrane, and peptide chain elongation (Figure 2h). In contrast, RK-33 treatment of SARS-CoV-2 infected cells results in a shift of pathways being impacted, with the five most significant pathways enriched being Mitochondrial translational initiation, Mitochondrial translational elongation, Mitochondrial translational termination, Antigen Presentation: Folding, assembly and peptide loading of class I MHC, and SRP-dependent cotranslational protein targeting to membrane (Figure 2i). We also queried the STRING database ^47^ and found pathways in RK-33 treated cells that were positively enriched (Endosomal/vacuolar pathway and Regulation of Complement cascade) and negatively enriched (Mitochondrial translation and Mitochondrial translation initiation). Pathways in virus infected DMSO control cells that were positively enriched included Regulation of IFNA signaling and Interferon alpha/beta signaling while Translation of Replicase and replication transcription complex and Endosomal sorting complex required for transport were negatively enriched (Figure S8). These data indicate that RK-33 treatment suppresses viral protein production corresponding to changes in the host proteomic landscape especially as it relates to innate immune response pathways.

### RK-33 inhibits HCoV-OC43 and SARS-CoV-2 production

Given that DDX3 has been shown to play a regulatory role in the viral biogenesis, we initially tested the impact of RK-33 on coronavirus replication using human coronavirus OC43 (betacoronavirus) as a BSL-2 model of SARS-CoV-2. The CC50 value of RK-33 on Rhabdomyosarcoma (RD) muscle cells (highly susceptible to HCoV-OC43 infection), was determined to be 3.22 μM (Figure 3a). Next the antiviral impact of RK-33 was tested on RD cells using non-toxic concentrations of RK-33 (0.1 and 1 μM). Treatment with 1 µM of RK-33 decreased HCoV-OC43 infectious titers by over 100-fold (>∼2 logs) (Figure 3b). As can be seen in Figure 3c, HCoV-OC43 infection of RD cells results in extensive cytopathic effect (CPE) by 4 dpi (left panel) that was partially relieved by 1 μM of RK-33 treatment (right panel). In addition, treatment with 1 µM of RK-33 decreased HCoV-OC43 infectious titers by over 100-fold (>∼2 logs) (Figure 3b). These data indicate that RK-33 is effective against a betacoronavirus, which encouraged us to test its impact on SARS-CoV-2.

**Figure 3.**
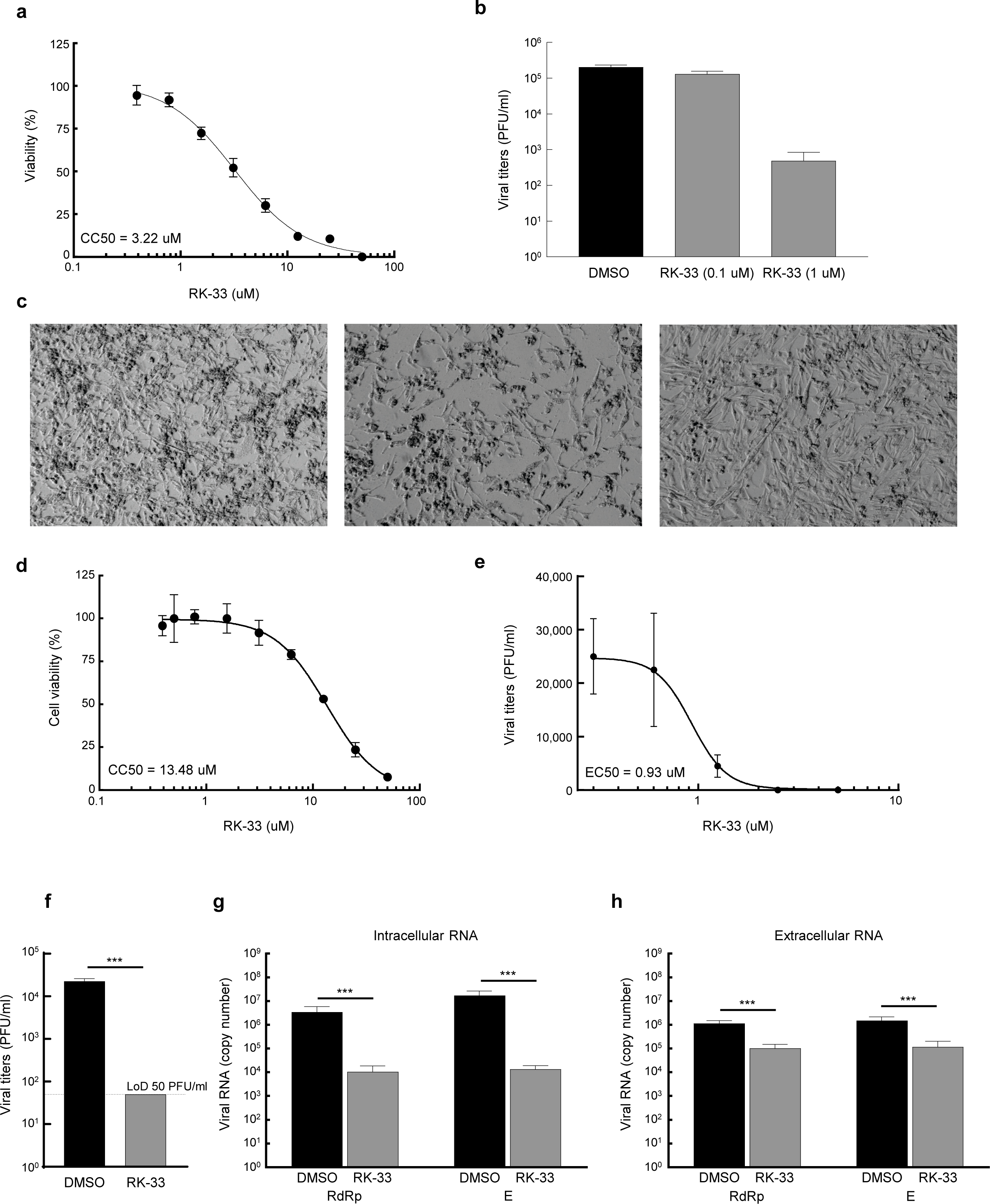
RK-33 decreases production of hCoV-OC43 and SARS-CoV-2. **a.** RD cells were treated with various concentrations of RK-33 and cell viability determined post-treatment using CellTiter Glo assay. **b.** RD cells were treated with RK-33 for 1 h followed by infection with hCoV-OC43 (MOI 0.1) and infectious titers at 3dpi were determined via plaque assay. **c.** RD cells were treated with RK-33 for 1 h followed by infection with hCoV-OC43 (MOI 0.1). CPE was observed at 4 dpi. **d.** Calu-3 cells were treated with various concentrations of RK-33 and cell viability determined using CellTiter Glo assay. **e.** Calu-3 cells were treated with various concentrations of RK-33 for 24 h followed by infection with SARS-CoV-2 Washington Variant (Lineage A) (MOI 0.1) in the presence of RK-33. Cells were also post-treated with RK-33. Supernatants were collected at 48 hpi and plaque assays performed to determine infectious titers. DMSO was included as a negative control. **f.** Calu-3 cells were treated with RK-33 (5 μM) or DMSO as described in panel E and viral titers determined by plaque assay. **g, h.** Intra and extra cellular SARS-CoV-2 RNA viral copy numbers as measured by RT-qPCR of RdRp and E genes. Data represents the average of 3 biological replicates and error bars display standard deviations. ***p-value<0.001.

Thus, since in our study we used Calu-3 cells (lung cancer cell line) as a model cell line for SARS-CoV-2 infection, our initial step in antiviral testing was to establish the CC50 of RK-33 in Calu-3 cells. For this purpose, cells were incubated with RK-33 in concentrations ranging from 0.39 to 50 µM for 24 h. Cell viability was measured by CellTiter-Glo assay which measures ATP production. The established CC50 for Calu-3 cell lines was 13.48 µM (Figure 3d). Next, the EC50 of RK-33 against SARS-CoV-2 was determined by treatment of Calu-3 cells with 0.3125 - 5 µM RK-33 or 0.05% DMSO for 24 h prior to infection. RK-33 and DMSO treated cells were infected with SARS-CoV-2 Washington isolate (Lineage A) at an MOI of 0.1 and 48 hpi the amount of infectious virus in the supernatants was determined by plaque assay. Overall, RK-33 displayed potent anti SARS-CoV-2 effect on Calu-3 cells with an EC_50_<1 µM (Figure 3e), resulting in a selectivity index (CC50/EC50) of 14.5. To confirm these results, a similar experiment was performed with cells treated with DMSO or 5 µM RK-33. In agreement with the results in Figure 3e, RK-33 treatment had a significant three log reduction in SARS-CoV-2 viral titers (Figure 3f), with no plaques detected in the RK-33 treated samples. Likewise, RK-33 treatment also resulted in a reduction of ∼2.5-3 log10 of intracellular viral RNA production as measured by RT-qPCR analysis using primers recognizing RdRp and E genes/regions (Figure 3g). Finally, a one log reduction in extracellular viral RNA (from cell-free supernatants) was also observed (Figure 3h) in RK-33 treated samples.

### Mutation status of SARS-CoV-2 proteins

Along with its worldwide spread, SARS-CoV-2 virus has mutated resulting in genome-wide amino acid variations across the four variants of varying lineages as displayed in Figure 4a. Protein N in the Alpha and Delta variants shows greater divergence from Lineage A than the spike proteins of these same variants. Proteins E, nsp7, and ORF3a have diverged more than the spike in the Beta variant, and the Delta variant displays some level of divergence from Lineage A in the M, nsp2, nsp4, nsp6, and NendU proteins. Due to large deletions and a frameshift, the ORF8 protein in the Delta variant has diverged from Lineage A to a greater extent than spike protein (https://covariants.org/ and Supp. Table 1). Furthermore, Figure 4b displays amino acid substitutions and deletions with respect to the Lineage A variant in sixteen of the proteins. This includes large truncations in ORF7a (Beta and Delta variants) and ORF8 (Alpha variant) and a high number of variations in the hACE2 receptor-interfacing spike proteins. Spike protein amino acid substitutions and deletions with respect to Lineage A variant are further visualized along the protein sequence at higher resolution in Figure 4c. These variations occur throughout the spike protein molecular structure, including within the S1-receptor-binding-domain as seen in the 3D models in Figure 4d. Details of mutations can be found in Supp. Table 1.

**Figure 4.**
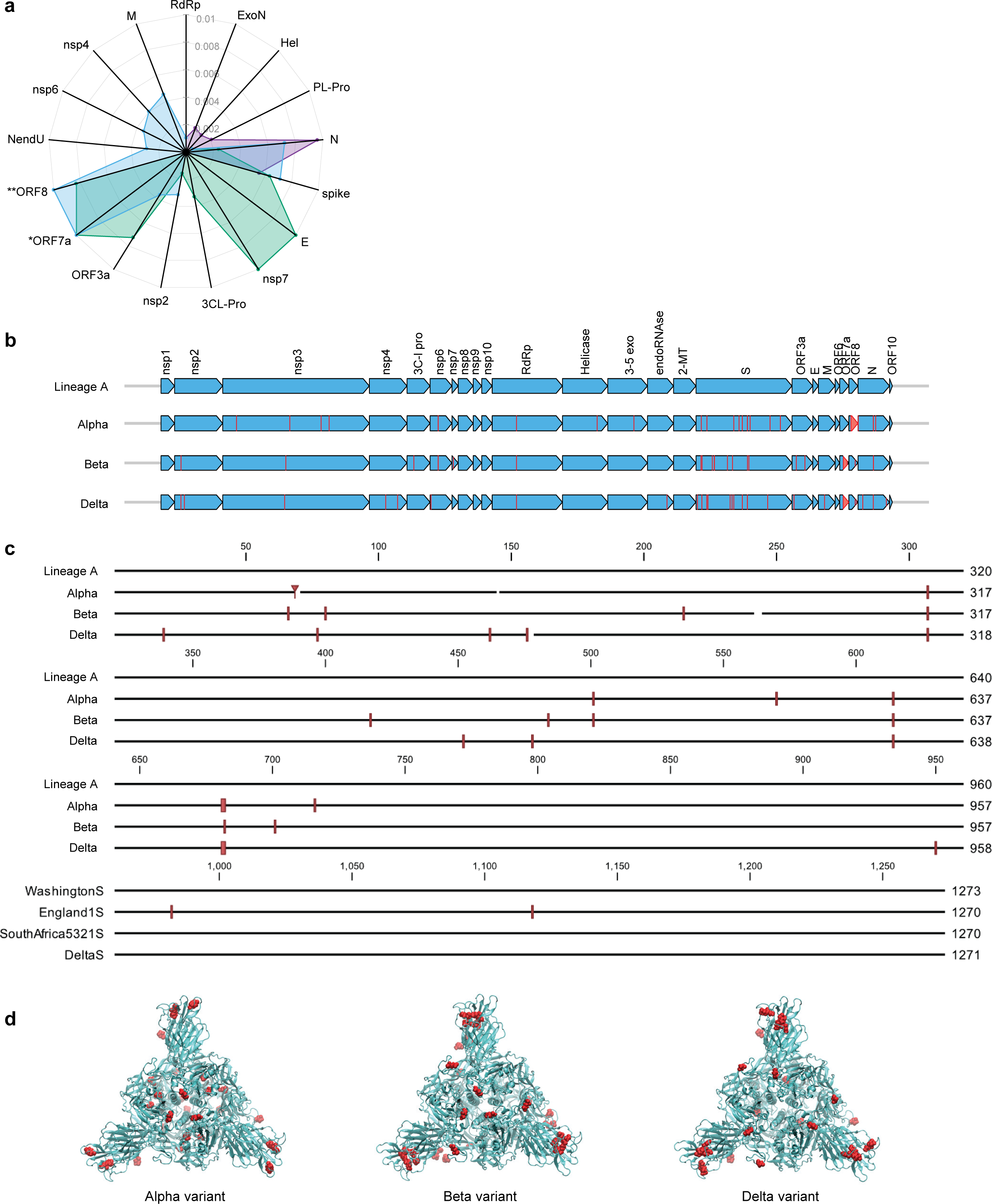
Comparison of mutation status of SARS-CoV-2 variants. **a.** Radar plot displaying estimated protein sequence evolutionary distance to the Lineage A variant for the Alpha (purple), Beta (green), and Delta (blue) variants. Only proteins that contain substitutions in at least one of the variants are shown. Proteins are ordered to group similar distances together for each isolate. *Delta ORF7a protein sequence has an estimated distance of 0.67 from Lineage A due to a deletion resulting in a frameshift. **Delta ORF8 protein sequence has an estimated distance of 0.02 from the Lineage A variant. These distances are plotted at the limits of the plot for clarity. **b.** Comparison of the SARS-CoV-2 proteome across the four variants of varying lineage used in this study. Arrows indicate all proteins encoded by the SARS-CoV-2 genome, and vertical lines denote amino acid variations with respect to the Lineage A variant. **c.** Multiple alignment of SARS-CoV-2 spike protein sequences for the four variants shown in b displayed in greater detail. Amino acid variations with respect to the Lineage A variant are indicated with red vertical lines. **d.** Spike protein homotrimer homology model of Lineage A variant with amino acid variations colored red for the Alpha, Beta, and Delta variants.

### RK-33 inhibits Alpha, Beta, and Delta variants of SARS-CoV-2

Due to the diversity between different viral variants, we next tested the effect of RK-33 on replication and virion production of different of SARS-CoV-2 variants. RK-33 treatment of cells resulted in a significant reduction (3-4 log fold) of infectious titers of the Alpha, Beta, and Delta variants (Figure 5a). Analysis of the intracellular and extracellular RNA (RdRp and E genes) also showed significant difference between RK-33 treated and DMSO control samples in the amount of the viral RNA production (Figure 5b-g). In conclusion, RK-33 displayed potent antagonistic SARS-CoV-2 effects against all tested variants.

**Figure 5.**
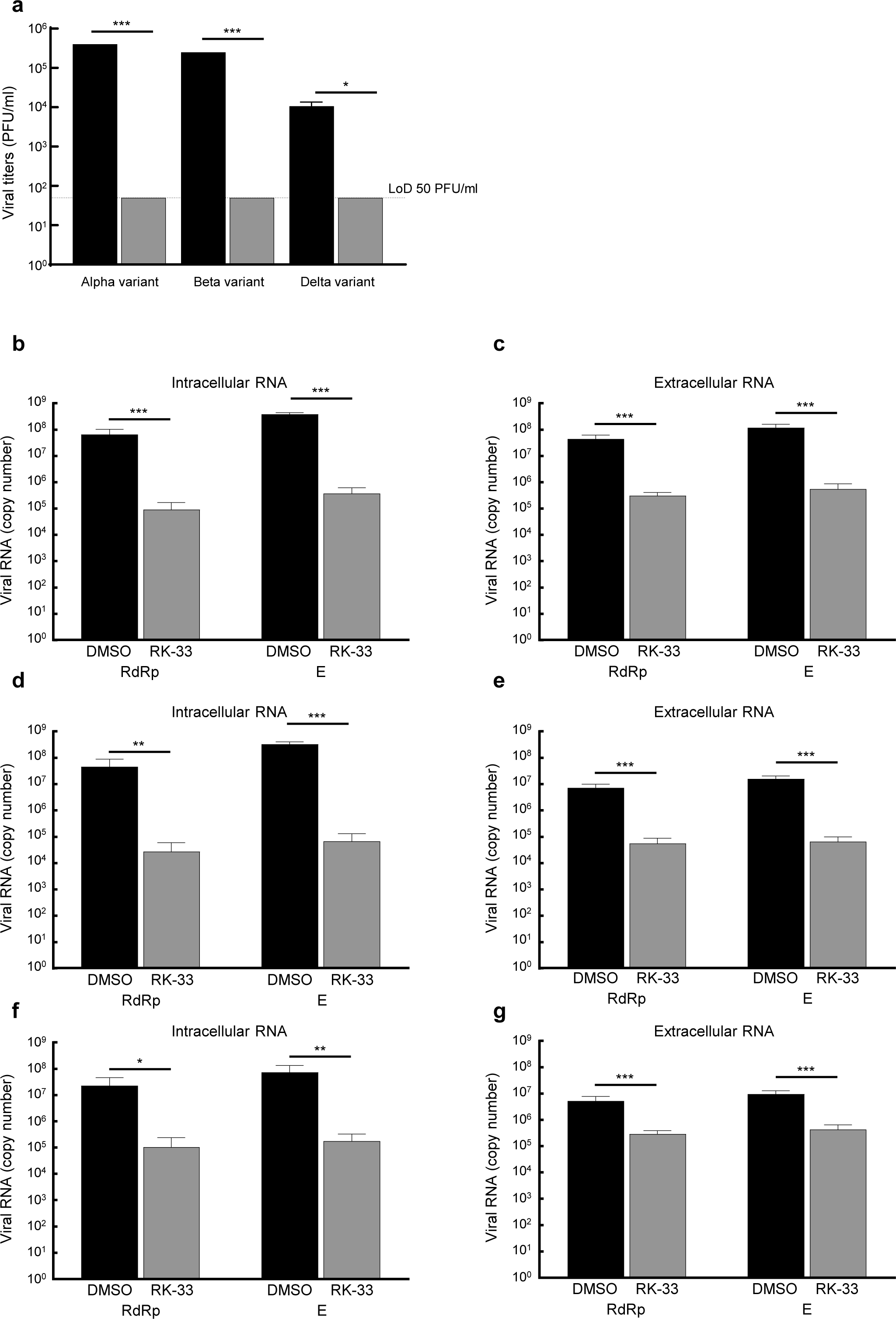
**a.** Calu-3 cells were treated with RK-33 (5uM) for 24 h followed by infection with Alpha, Beta, and Delta variants of SARS-CoV-2. Supernatants were collected at 48 hpi and plaque assays performed to determine infectious titers. DMSO was included as a negative control. **b.** RNA was extracted from SARS-CoV-2 Alpha variant-infected cells treated with RK-33 and subjected to RT-qPCR in a one-step protocol that amplified RdRp and E genes. Copy numbers were calculated from known positive controls. **c.** Extracellular RNA was extracted from above cell supernatants and processed as earlier. **d, e.** Results from intra- and extracellular RNA from RK-33 treatment of Beta variant infection. **f, g.** Results from intra and extracellular RNA from RK-33 treatment of Delta variant.

### RK-33 effects TMPRSS2 and reduces Spike-pseudotyped lentivirus constructs in vitro

The transmembrane serine protease TMPRSS2 has been identified as the protease that cleaves spike protein of SARS-CoV-2 thus facilitating entry ^30^. To determine if abrogating DDX3 activity by RK-33 would affect TMPRSS2 expression, we treated Calu-3 cells with different concentrations of RK-33 for 48 h and probed for the levels of TMPRSS2 and DDX3 by immunoblotting. As shown in Figure 6a, RK-33 at 7.5 µM (CC25) was able to reduce TMPRSS2 expression by 60% percent, quantified using Fiji (ImageJ). To determine if the decrease in TMPRSS2 level by RK-33 had any biological significance on viral entry, we treated Calu-3 cells with RK-33 for 24 h, followed by infection with a SARS-CoV-2 Spike-pseudotyped GFP lentivirus construct.

**Figure 6.**
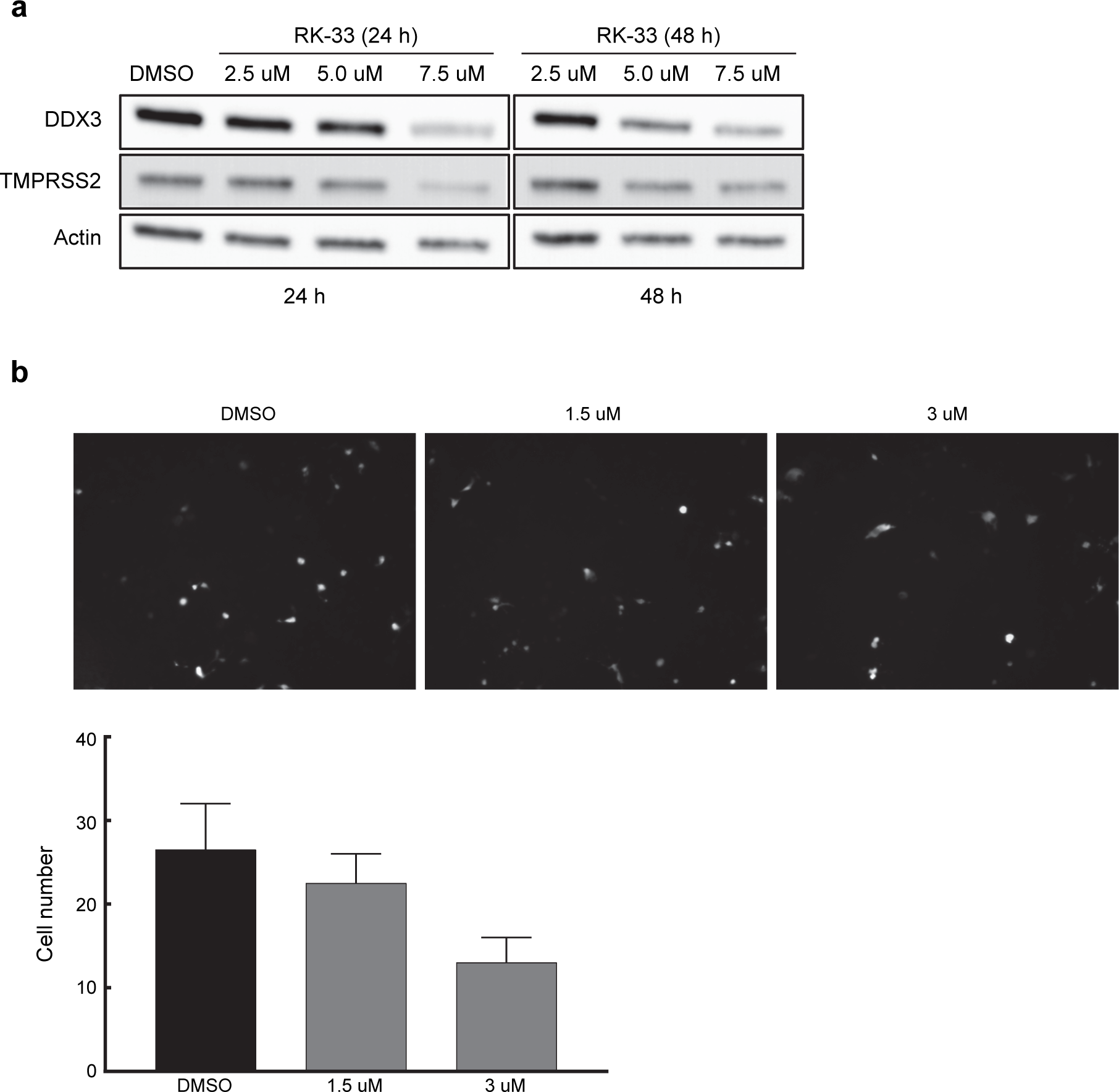
**a.** Immunoblot of proteins extracted from increasing concentrations of RK-33 treatment of Calu-3 cells for 24 and 48 h. **b.** Effect of increasing concentrations of RK-33 on Calu-3 cells for 48 h that were infected with pseudotyped virus. **c.** Quantification of Calu-3 cells seen in b.

Two days following infection, we counted the GFP cells using a random unbiased approach. As shown in Figure 6b, use of RK-33 at CC5 (3 µM), reduced the infected cell numbers by 50%. This supports our hypothesis that dysregulating DDX3 by RK-33 can affect TMPRSS2 expression resulting in decrease of SARS-CoV-2 lentivirus infection of Calu-3 cells.

## Discussion

SARS-CoV-2 has been successfully targeted in a few ways. Prophylactic vaccines are available that reduce/eliminate hospitalization and associated medical complications due to COVID-19. Presently, other treatments are limited to the antiviral nucleoside analog Remdesivir from Gilead Biosciences and a monoclonal antibody cocktail from Regeneron. However, Remdesivir is only prescribed in severe cases, and both require intravenous infusion making it difficult to obtain without hospitalization. Moreover, Remdesivir has shown mixed results in reducing the time to recovery in hospitalized patients ^48–52^. Another nucleoside analog drug Molnupiravir (EIDD-2801) from Merck is in the process of FDA approval which reduces the risk of hospital admission or death by approximately 50% in patients at risk for poor outcomes ^53^.

On the other hand, identifying virus-host interacting factors could reveal new targets to mitigate medical complications associated with SARS-CoV-2 infection. Over the years, several roles of a host protein, DDX3, have been identified as crucial for replication and production of many viruses^54, 55^. Interestingly, DDX3 has also been identified, using proteomics, to be involved in SARS-CoV-2 replication ^54, 55^. Moreover, computational and system biology approaches have indicated different proteins of SARS-CoV-2 to be associated with cellular organelle functions that are regulated by DDX3 activity ^56^. In this study, we used a small molecule inhibitor of DDX3, RK-33, to study its impact of SARS-CoV-2. Using the Lineage A variant, we showed that RK-33 reduced viral titers by over 10,000-fold by plaque assay. These data were further validated by RT-qPCR (RdRp and envelope) of the intra- and extracellular SARS-CoV-2 RNA of the RK-33 treated Lineage A infected Calu-3 cells. This work is in agreement with a study showing that inhibition of DDX3 by two different DDX3 inhibitors, RK-33 and C-4B, impacted SARS-CoV-2 RNA production, but this study did not determine the impact on viral titers ^54^.

Based on this encouraging finding, we performed RNA-seq and proteomics analysis on SARS-CoV-2 infected Calu-3 cells, before and after RK-33 treatment. PCA of the RNA showed the major effect of RK-33 on the cells. It was interesting to note that in the virus infected Calu-3 cells, more than 80% of the RNA species was that of SARS-CoV-2, while in the RK-33 group, the host mRNA species was 80% represented. This is indicative that perturbing DDX3 by RK-33 restored many of the host cellular functions while suppressing SARS-CoV-2 replication. This could be clearly seen in both the heat map and the PCA plot with RK-33 treatment serving to “normalize” expression of the virus infected cells towards the uninfected cell type gene expression. Volcano plots also demonstrated how the high gene expression patterns caused by the virus infection were modulated by RK-33 treatment of the cells. Given that the sensitivity of RK-33 to reduce Lineage A titers is indicative of the potential of targeting host proteins for SARS-CoV-2 treatment.

To further characterize the effect of RK-33 on the host cellular machineries, we performed proteomics on SARS-CoV-2 infected Calu-3 cells. Proteomic-based PCA analysis showed that RK-33 treated virus infected samples segregated from the untreated infected samples. The results indicated that RK-33 was the major determinant of segregation in the PCA plot. Subsequent pathway analysis showed that many of the downregulated pathways were associated with virus attachment, entry, assemble and release. Proteomics analysis clearly demonstrated that most of the proteins involved in SARS-CoV-2 genome replication and transcription (ORF1 ab, ORF3A, Spike, Membrane, ORF7A and Nucleoprotein) were significantly downregulated by RK-33 in SARS-CoV-2 infected Calu-3 cells. The ability of RK-33 to reduce viral titers of SARS-CoV-2 Lineage A could reflect on the dependence of SARS-CoV-2 on DDX3 for its replication. This is probably because at the molecular level SARS-CoV-2 usurps DDX3 in the host cell to its advantage for replication and for pathogenesis. This is in parallel to what is already demonstrated by many other viruses that utilize DDX3 for viral entry, processing, and egress from the host cells ^57^. Furthermore, as DDX3 has been shown to interact with nucleocapsid protein of SARS-CoV-2 ^54^, it is plausible that the use of RK-33 is directly interfering with SARS-CoV-2 replication machinery as well.

Based on these promising results, we evaluated the effect of RK-33 on viral titers in the Alpha, Beta, and Delta variant infected cells. RK-33 was able to reduce the viral loads by 200,000, 125,000, and 2,500-fold respectively. This is not unexpected as targeting host factors essential for viral replication, such as DDX3, would have similar outcomes for all the variants. As is evident from the spike protein modeling, there is a higher concentration of spike protein mutation within the ACE-2 binding region, but this does not affect the ability of RK-33 to reduce viral titers across all the variants. Also, the amino acid substitutions lead to increased evolutionary distances in different proteins among the four isolates used in this study. Thus, the four variants used in this study could serve as diverse models adapted to provide a growth advantage or independent evolution of SARS-CoV-2 leading to increased virulence and pathogenesis.

Furthermore, we showed that RK-33 was able to reduce TMPRSS2 expression in Calu-3 cells. A key aspect of the TMPRSS2 gene is the presence of guanine-rich tracts of DNA in its gene promoter that can form G-quadruplex structures. These structures can be targeted by benzoselenoxanthene derivatives that stabilize TMPRSS2 G-quadruplexes in vitro, thus downregulating TMPRSS2 gene expression ^58^. In this context, DDX3 was found to facilitate both translation of general complex secondary structures including G-quadruplexes ^59, 60^, as well as mRNAs with secondary structure in immediate vicinity to their m^7^GTP cap ^61^. We have presented data that demonstrates that use of RK-33 even at CC5, reduced the SARS-CoV-2 infection rate by 50%.

These data indicate that targeting DDX3 by RK-33 for SARS-CoV-2 treatment could potentially reduce SARS-CoV-2 virulence and pathogenicity by two different mechanisms. First, it can decrease the infection rate by downregulating TMPRSS2 protein expression, and second, reduce the pathogenicity, by suppressing SARS-CoV-2 replication. This two-prong mechanism of action would be extremely effective at suppressing viral replication of any newly emerging variants of concern, which is in line with an antiviral that is targeting a host factor. We fully appreciate that the impact of RK-33 on TMRPSS2 expression is only one mechanism by which RK-33 may be impacting SARS-CoV-2 production. Indeed, our proteomic and transcriptomic analysis points to multiple cellular events being altered in infected cells by RK-33 treatment, such as innate immune responses, translation, and membrane trafficking, each of which are important for viral production.

In conclusion, we report on the importance of DDX3 and its targeting by the small molecule inhibitor RK-33 which was developed earlier by rational drug design that resulted in a unique structural scaffold with little or no toxicity in animals ^7–9, 62^. Use of RK-33 on Calu-3 cells resulted in suppressing virus titers from all four isolates and has the potential to treat SARS-CoV-2 infection. We are the first to present a promising HTA strategy to tackle the different variants associated with SARS-CoV-2 as well as subsequent variants that may arise.

### Methods Virus inhibitor

RK-33 was dissolved in 100% DMSO to the final 10 mM concentration. For a cell-based assay the stock was diluted in complete cell media to required concentrations.

### Cell culture and viruses

Calu-3 (ATCC HTB-55) and Vero (ATCC CCL-81) cells were cultured at 37°C and 5%CO2 in EMEM and DMEM, respectively, supplemented with 10% FBS, 1% L-glutamine, and 1% penicillin-streptomycin. The following SARS CoV-2 isolates, obtained from BEI Resources, were used in the study: Lineage A (NR-52281), Beta variant (NR-54008), Alpha variant (NR-54000), Delta variant (NR-55611).

### CC_50_ and EC_50_

For CC_50_ assays cells were maintained as described above in 96-well white well plates for 24 h prior to RK-33 drug treatment. Cells (2.8 x 10^4^) were incubated with RK-33 in serially diluted concentrations from 50 to 0.39 µM for 24 h. Cell viability was estimated by CellTiter-Glo assay (Promega) in accordance with manufacturer’s recommendation. For EC_50_ assays Calu-3 cells were maintained as described above in 12-well plates for 24 h prior to EC_50_ assays. Cells were treated with RK-33 in decreasing concentrations from 5µM to 0.3 µM for 1 h followed by one h of SARS CoV-2 Lineage A variant infection at multiplicity of infection (MOI) 0.1. After one h of incubation, viral infectant was removed, cells were washed one time with PBS and fresh complete medium supplemented with RK-33 in equivalent concentrations was added. Virus containing cell media was collected 48 hpi and virus titers were estimated by standard plaque assay as described elsewhere ^63, 64^.

### Cell infection

Calu-3 cells (8 x 10^5^) were plated in 12-well plates 24 h prior to drug treatment. Cells were then incubated with RK-33 at 5µM concentration for 24 h followed by infection with SARS CoV-2 variant as indicated, at MOI 0.1. One hour post infection infectant was removed, cells were washed one time with PBS and fresh complete medium supplemented with RK-33 at 5µM concentration was added. Cells infected with SARS CoV-2 as well as virus containing supernatants were collected 48 h post infection.

### RNA-Seq

Total RNA was extracted from virus containing cell lysates using Trizol LS (Invitrogen, Carlsbad, CA) and purified using a column purification method (Qiagen, Germany) in accordance with manufacturer’s recommendations that were modified to isolate both small and large RNA species. RNA concentration and preliminary quality control was estimated on a NanoDrop spectrophotometer (ThermoFisher Scientific) while RNA-Seq was performed commercially (BGI Americas, San Jose, CA) as detailed in Supplementary Methods. Four biological replicates were analyzed from each of the treatments (Uninfected cells treated with DMSO, Uninfected cells treated with RK-33, Infected cells treated with DMSO, and Infected cells treated with RK-33).

### Proteomics

Proteomics was performed at the Johns Hopkins University School of Medicine Mass Spectrometry and Proteomics Core using 50 ug of each sample as detailed in Supplementary Methods. Biological replicates of four were analyzed as in RNA-Seq.

### RT-qPCR and RNA isolation

Viral RNA was extracted from virus containing cell media and from cell lysates using Trizol LS (from Invitrogen, cat# 10296010) and purified by Direct-Zol RNA purification kit (from Zymo research, cat#R2052) in accordance with manufacturer’s recommendations. SARS-CoV-2 RNA (RdRp and E genes) was amplified by the one-step Logix Smart-2 COVID-19 real-time PCR reagents kit from Co-Diagnostics, Inc. (Salt Lake City, UT, USA).

Normalization was performed using RNaseP gene amplified from host RNA. Copy numbers were estimated based on the included quantified positive control reagent.

### Mutational analysis

The genome sequence of the Lineage A variant was retrieved from GenBank accession MW811435, a direct submission of NR-52281 sequencing at ATCC. The Alpha variant genome sequence was retrieved from GenBank accession MZ376737. The Beta variant genome sequence was retrieved from GenBank accession MZ345001. The Delta variant genome sequence was reconstructed from GenBank accession MN908947 with variations as reported on the NR-55611 Certificate of Analysis. Of particular note is a deletion within ORF7a (AA 44-100) with a frameshift that results in extension of the protein sequence with a new stop codon downstream of ORF7b.

Protein sequences for 25 SARS-CoV-2 proteins were extracted from the four genome sequences. For each protein, a four-isolate multiple sequence alignment was performed in CLC Genomics Workbench with a progressive alignment algorithm using default parameters. Differences with respect to the Lineage A variant were noted. Evolutionary distances of proteins were estimated using the WAG substitution model ^67^.

Three-dimensional structural models of the reference A lineage spike protein homotrimer in open and closed conformations were obtained from the Zhang Lab (zhanggroup.org/COVID-19). This model was generated using the D-I-TASSER/ C-I-TASSER pipeline ^68^. The protein sequence of the structural model is identical to that of the Lineage A variant used in this study. Residues with variations found in the Alpha, Beta and delta variants were visualized on this structure with the Visual Molecular Dynamics program ^69^.

## Acknowledgements

The following SARS-CoV-2 isolates were obtained through BEI Resources, NIAID, NIH: SARS-Related Coronavirus 2, Isolate USA-WA1/2020, NR-52281, SARS-Related Coronavirus 2, Isolate hCoV-19/South Africa/KRISP-EC-K005321/2020, NR-54008, contributed by Alex Sigal and Tulio de Oliveira, SARS-Related Coronavirus 2, Isolate hCoV-19/England/204820464/2020, NR-54000, contributed by Bassam Hallis, SARS-Related Coronavirus 2, Isolate hCoV-19/USA/PHC658/2021 (Lineage B.1.617.2; Delta Variant), NR-55611, contributed by Dr. Richard Webby and Dr. Anami Patel.

Pseudotyped viruses were a kind gift from Dr. Prashant Desai (JHU SKCCC).

We wish to acknowledge Robert O’Malley and Bob Cole at the Johns Hopkins University School of Medicine Mass Spectrometry and Proteomics Core for performing the proteomics experiments.

We wish to acknowledge Co-Diagnostics, Inc. (Salt Lake City, UT, USA) for providing Logix Smart COVID-19 real-time PCR reagents.

We wish to acknowledge the NIH (Grant R01CA207208) and FAMRI for funding to VR. Support was also provided by Drs. Bhujwalla and Horton and the Department of Radiology.

## Author contributions

FV, IA, KH, VR planned the experiments. FV, IA, PW, and SL performed the experiments. FV, IA, KH, VR analyzed the results. FV, IA, PW, KH, VR wrote the manuscript.

## Competing Interests statement

VR holds a patent on the composition of RK-33 (US patent # 8,518,901).

## Supplementary Methods

### RNA-Seq

Total RNA was checked for integrity on a bioanalyser and stranded libraries were constructed from oligo dT purified mRNAs. RNA was fragmented and first-strand cDNA was generated using random N6-primed reverse transcription, followed by a second-strand cDNA synthesis with dUTP instead of dTTP. The synthesized cDNA was subjected to end-repair, 3’ adenylated, and adaptors were ligated to cDNA fragments. The dUTP-marked strand was degraded by Uracil-DNA-Glycosylase (UDG) and the remaining strand was PCR amplified to generate a cDNA library. After heat denaturation of the library, the single strand DNA was cyclized by splint oligo and DNA ligase. This is followed by rolling circle replication and DNA nanoball synthesis and eventually sequencing on the DNBSEQ (DNBSEQ Technology) platform.

Quality control (QC) was then performed on the raw reads to determine whether the sequencing data is suitable for subsequent analysis. After QC, the filtered clean reads were aligned to the reference human genome (hg38) using HISAT2 and the statistics of the mapping rate and the distribution of reads on the reference sequence were used to determine whether the alignment result passes the second QC of alignment using FastQC. This was followed by gene quantification analysis and other analysis based on gene expression such as PCA, correlations, and differential gene screening using DeSEQ2. We also performed significant enrichment analysis of GO function on differentially expressed genes and significance enrichment analysis of pathways analysis using REACTOME.

### Proteomics

Proteins (50 ug) were reduced with 50mM Dithiothreitol in 10 mM Triethylammonium bicarbonate (TEAB) at 60C for 45 minutes followed by alkylating with 100 mM Iodoacetamide in 10 mM TEAB at room temperature in the dark for 15 minutes. MS interfering reagents were removed by precipitating 50 ug proteins by adding 8 volumes of 10% trichloroacetic acid in cold acetone at -20C for 2 h. The pellet was centrifuged at 16,000 g for 10 minutes at 4C. The TCA/Acetone supernatant was removed, and the protein pellet was washed with an equivalent 8 volumes acetone at -20C for 10 minutes prior to centrifuging at 16,000 g for 10 minutes at 4C. The acetone supernatant was removed from the protein pellet. The four sets of 16 protein pellets (50 ug) were resuspended and digested overnight at 37C in 100 uL 50 mM TEAB with 5 ug Trypsin/Lys-C per sample. Each sample was labeled with a unique TMTpro 16-plex reagent (Thermo Fisher, LOT # VJ313476) according to the manufacturer’s instructions and quenched with 5ul of 5% hydroxylamine for 15 minutes. All 16 TMT labeled peptide samples in each of the 4 sets were combined and dried by vacuum centrifugation. The combined TMT-labeled peptides (800ug) were re-constituted in 100 µL 200mM TEAB buffer and filtered through Pierce Detergent removal columns (Fisher Scientific PN 87777) to remove excess TMT label, small molecules and lipids. Peptides in the flow through were diluted to 2 mL in 10 mM TEAB in water and loaded on a XBridge C18 Guard Column (5 µm, 2.1 x 10 mm, Waters) at 250 µL/min for 8 min prior to fractionation on a XBridge C18 Column (5 µm, 2.1 x 100 mm column (Waters) using a 0 to 90% acetonitrile in 10 mM TEAB gradient over 85 min at 250 µL/min on an Agilent 1200 series capillary HPLC with a micro-fraction collector. Eighty-four 250 ul fractions were collected and concatenated into 24 fractions and dried ^65^. Peptides in each of the 24 fractions were analyzed on a nano-LC-Orbitrap-Fusion Lumos-IC in FTFT mode (Thermo Fisher Scientific) interfaced with an EasyLC1200 series by reversed-phase chromatography using a 2%–90% acetonitrile in 0.1% formic acid gradient over 90 minutes at 300 nl/min on a 75 µm x 150 mm ReproSIL-Pur-120-C18-AQ column 3 µm, 120 Å (Dr.Maisch). Eluting peptides were sprayed into the mass spectrometer through a 10 µm emitter tip (New Objective) at 2.6 kV. Survey scans of precursor ions were acquired from 350-1400 m/z at 120,000 resolution at 200 m/z. Precursor ions were individually isolated within 0.7 m/z by data dependent monitoring and 15s dynamic exclusion, and fragmented using an HCD activation collision energy 34 at 50,000 resolution. Fragmentation spectra were processed by Proteome Discoverer v2.4 (PD2.4, ThermoFisher Scientific) and searched with Mascot v.2.8.0 (Matrix Science, London, UK) against RefSeq2021_204 Human database. Search criteria included trypsin enzyme, one missed cleavage, 3 ppm precursor mass tolerance, 0.01 Da fragment mass tolerance, with TMTpro on N-terminus and carbamidomethylation on C as fixed and TMTpro on K, oxidation on M, deamidation on N or Q as variable modifications. Peptide identifications from the Mascot searches were processed within PD2.4 using Percolator at a 5% False Discovery Rate confidence threshold, based on an auto-concatenated decoy database search. Peptide spectral matches (PSMs) were filtered for Isolation Interference <30%. Relative protein abundances of identified proteins were determined in PD2.4 from the normalized median ratio of TMT reporter ions, having signal to noise ratios >1.5, from all PSMs from the same protein. Technical variation in ratios from our mass spectrometry analysis is less than 10% ^66^.

## Supplementary Figure Legends

**Figure S1.**
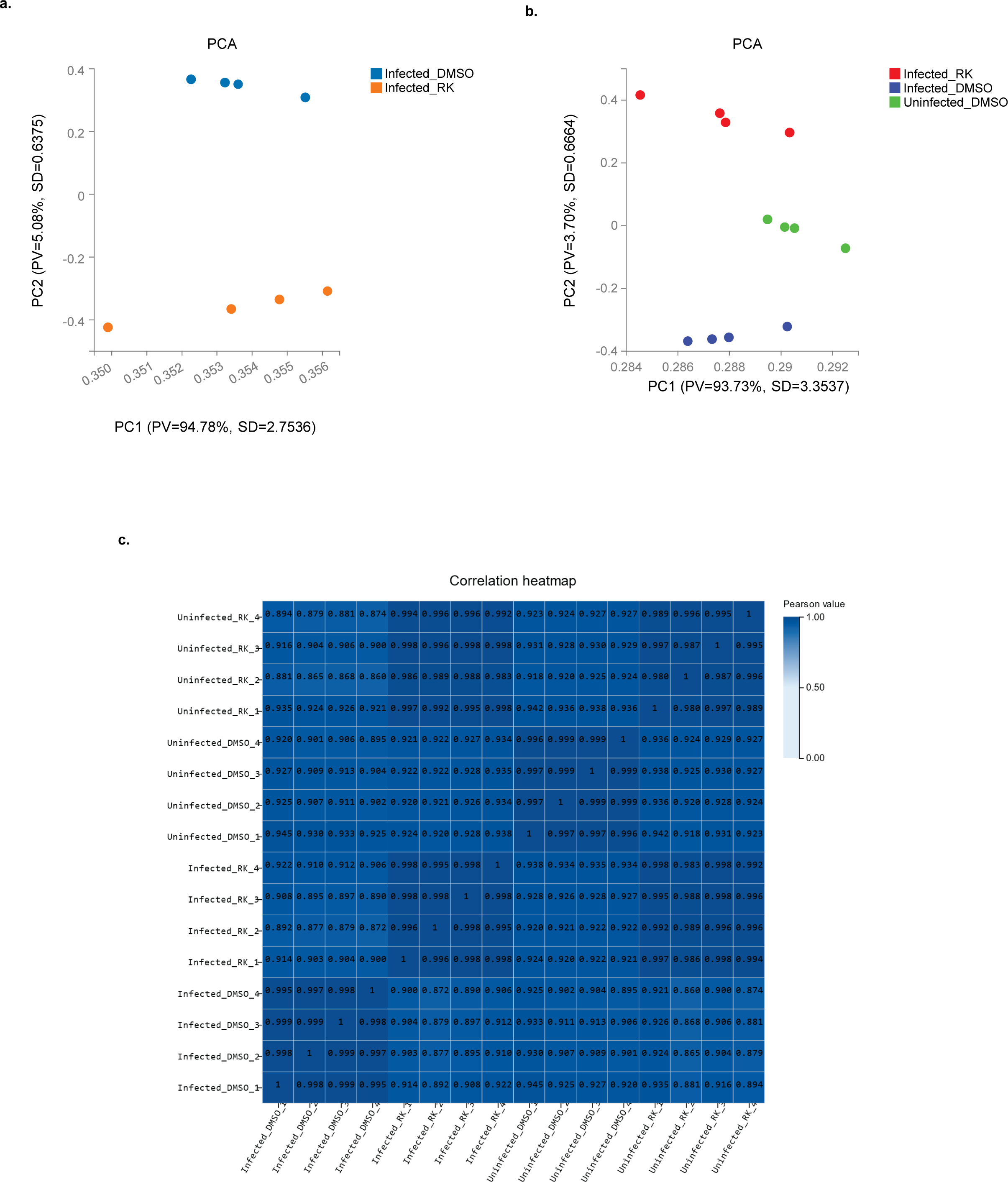
**a.** PCA plot of RK-33 treated virus infected samples (blue) and DMSO control treated virus infected samples (orange). PV means "Proportion of variance", SD means standard deviation. **b.** PCA plot of RK-33 treated virus infected samples (red), DMSO control treated virus infected samples (blue), and DMSO treated uninfected samples (green). PV means "Proportion of variance", SD means standard deviation. **c.** A correlation plot to study the correlation of gene expression between samples. Pearson correlation coefficients of all gene expression between every sample pair was calculated, and these coefficients were plotted in the form of a heatmap. The correlation coefficients demonstrate similarity of overall gene expression between each sample with higher correlation coefficient being more similar the gene expression level. Higher correlation coefficients are represented by darker colors while lighter colors represent lower correlations.

**Figure S2.**
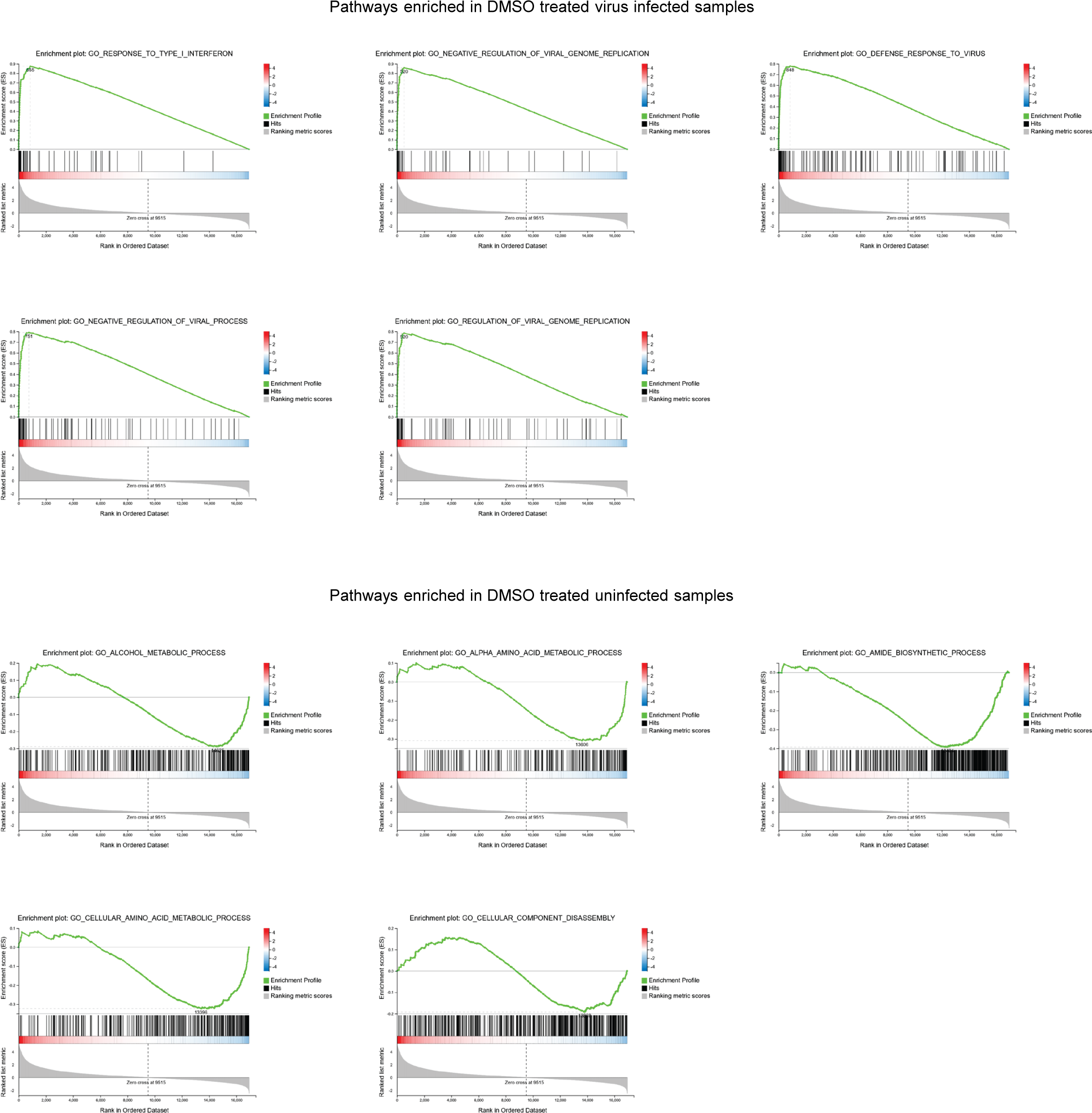
Enriched pathways obtained by GSEA of DMSO treated infected samples and DMSO treated uninfected samples. GSEA was performed using the Molecular Signatures Database collection Gene Ontology (GO) Biological Process. Displayed are the top five pathways enriched in DMSO treated samples and top five enriched in uninfected DMSO samples.

**Figure S3.**
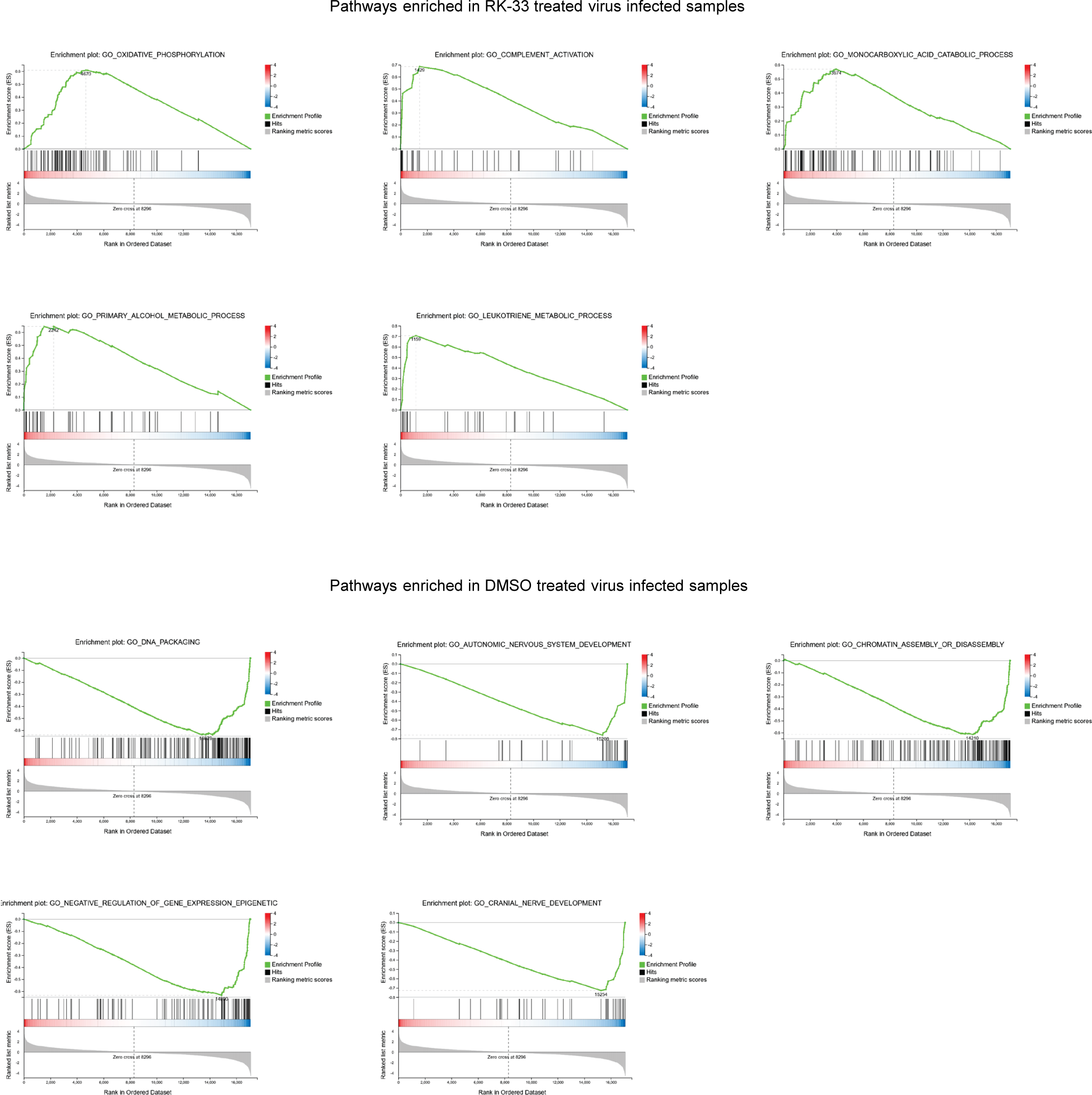
Enriched pathways obtained by GSEA of RK-33 treated virus infected samples and DMSO treated virus infected samples.

**Figure S4.**
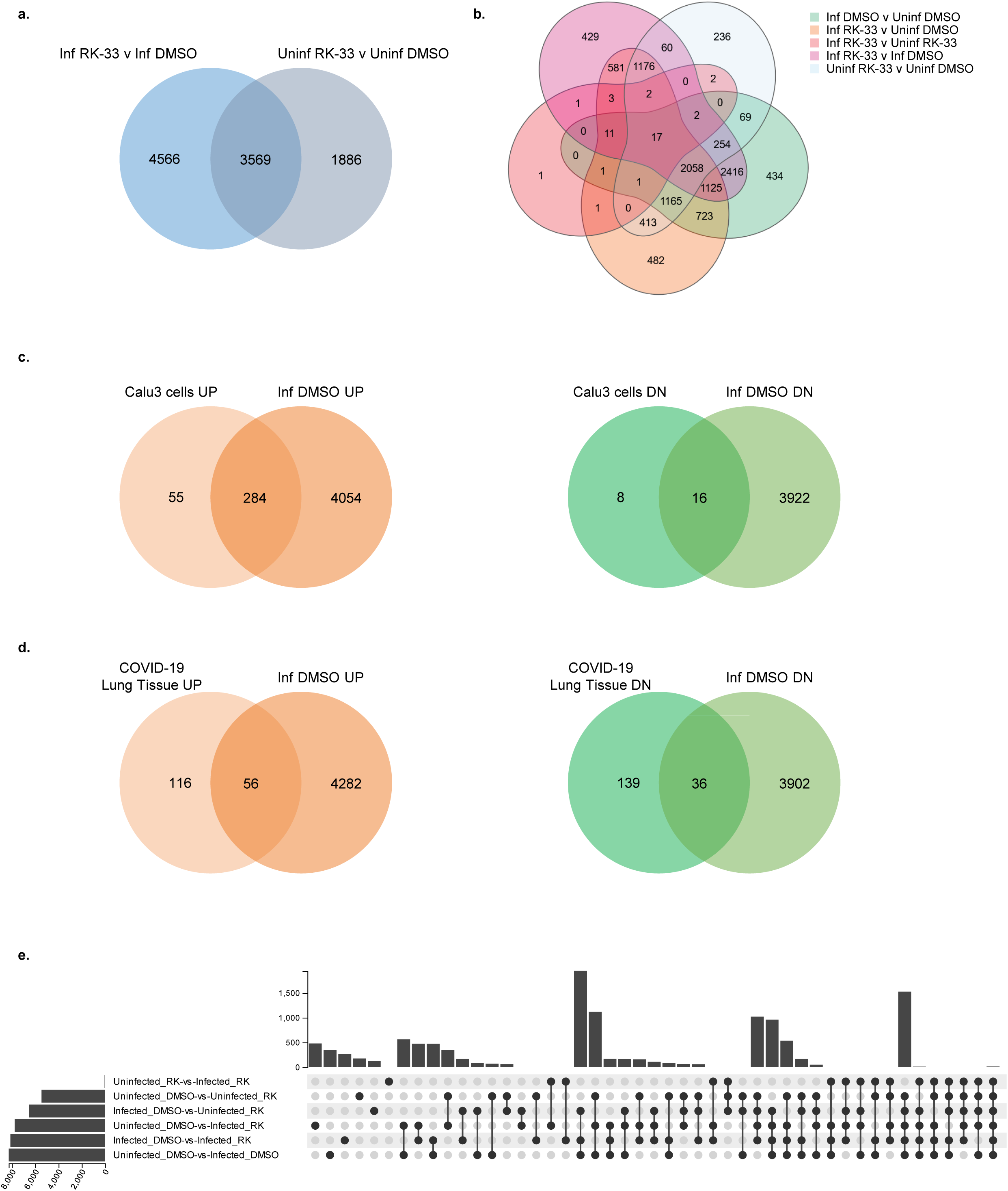
**a.** Venn diagram displaying gene sets dysregulated by RK-33 treatment of virus (Inf RK-33 v Inf DMSO) compared to genes dysregulated by RK-33 alone (Uninf RK-33 v Uninf DMSO). **b.** A Venn diagram displaying gene set comparison between all sample of all major comparisons. **c.** Venn diagrams comparing gene sets from this work with published data from Calu-3 cells. **d.** Venn diagrams comparing our gene sets with published work from COVID-19 patient lungs extracted post-mortem. **e.** Venn diagram of differentially expressed genes from RNA-Seq analyzed samples. The left bar graph displays the number of genes, and the Y-axis represents the name of gene set. In the upper right histogram, the X-axis displays the intersection of different gene sets, and the Y-axis shows the number of genes. Each column in the lower right shows the relationship between the left gene set and the upper intersection and the corresponding number of genes in common.

**Figure S5.**
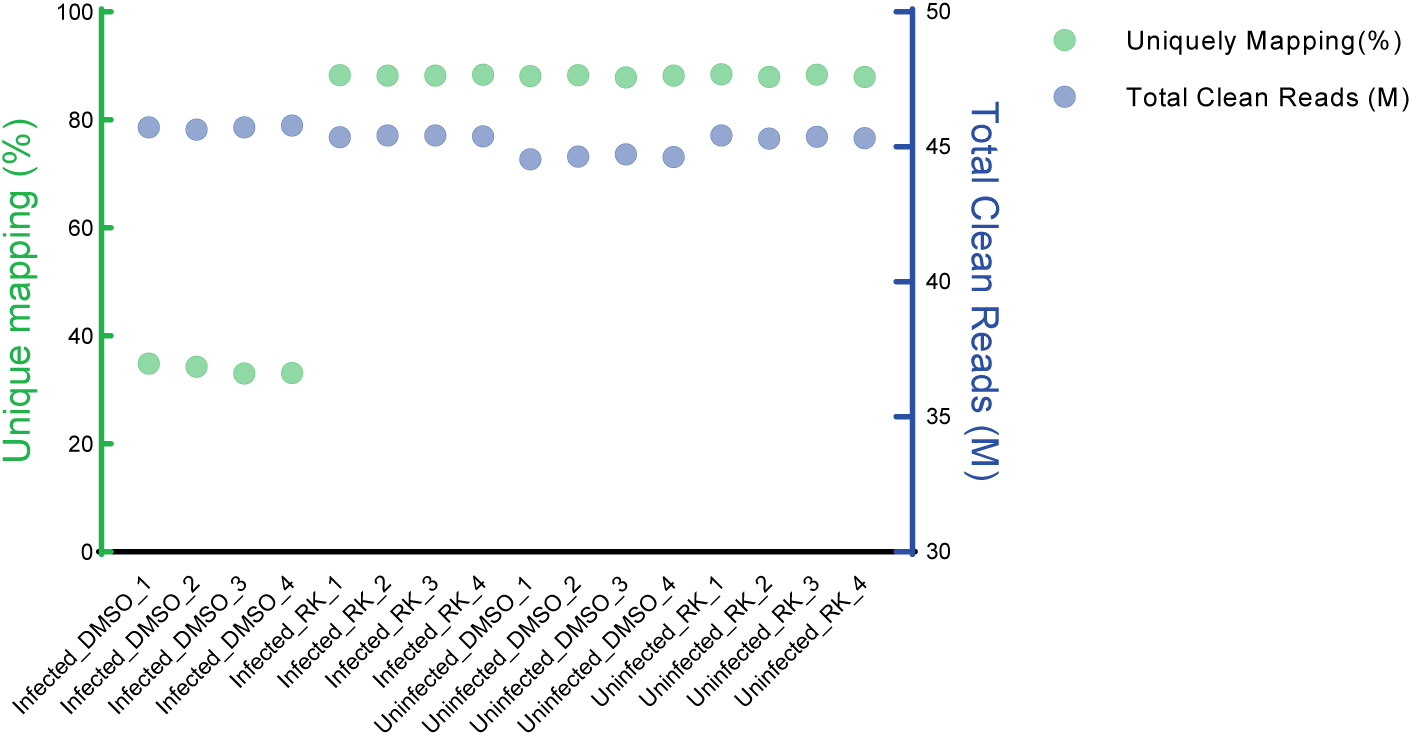
Scatterplot displaying the QC of the RNA-seq of all the samples mapped to the human genome. Left Y-axis displays the percent of uniquely mapping reads (green) and the right Y-axis displays total reads in millions (blue).

**Figure S6.**
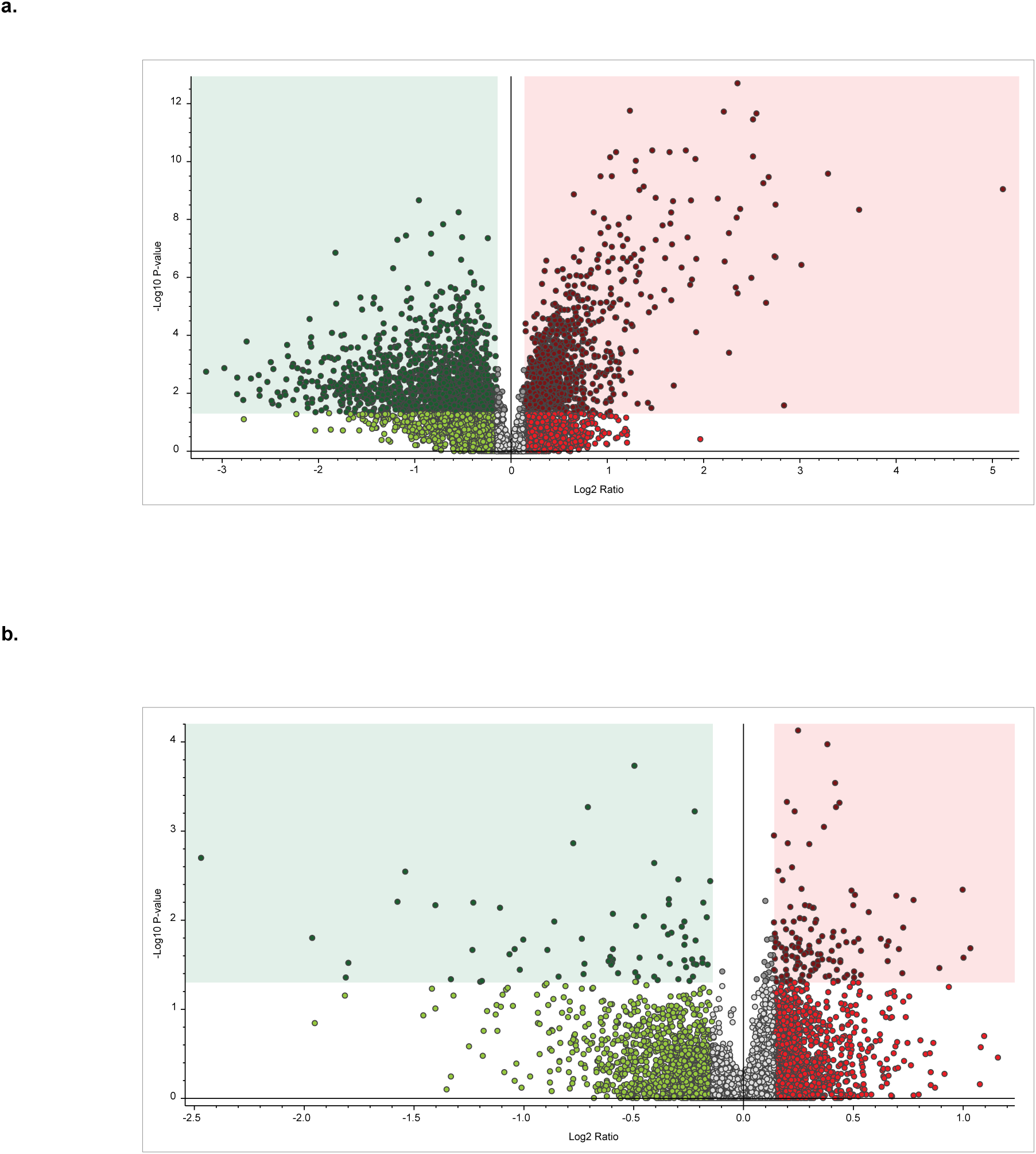
**a.** Volcano plot of significantly changed proteins (P<0.05, FC>1.1) in RK-33 treated uninfected samples compared to DMSO treated uninfected samples. **b.** Volcano plot of significantly changed proteins (P<0.05, FC >1.1) in RK-33 treated infected samples compared to RK-33 treated uninfected samples.

**Figure S7.**
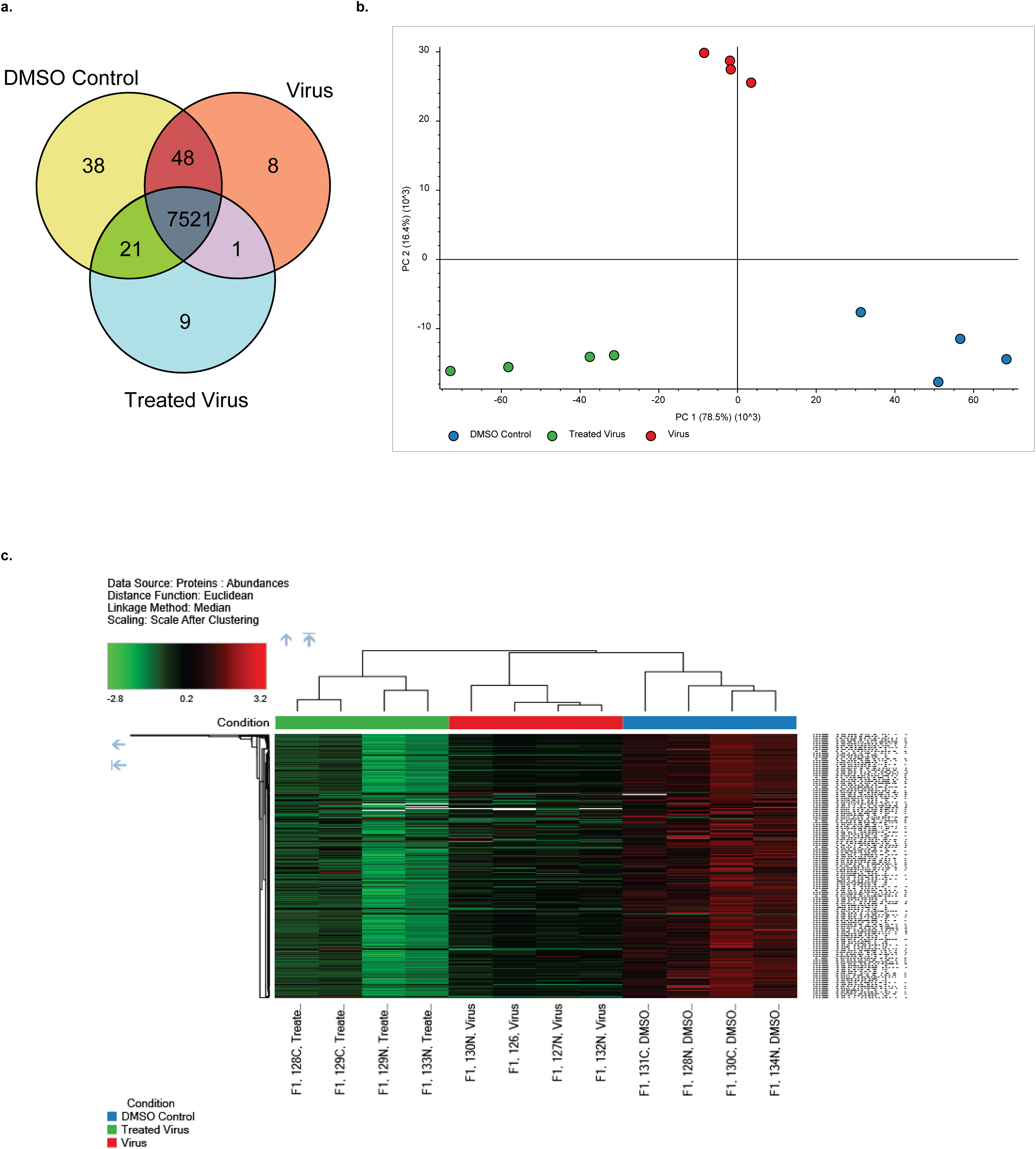
**a.** Venn diagram displaying proteins that are found in DMSO treated uninfected samples (DMSO control), DMSO treated infected samples (Virus), and RK-33 treated infected samples (Treated virus). **b.** PCA plot of proteomics samples - DMSO treated uninfected samples (DMSO control) (blue), DMSO treated infected samples (Virus) (red), and RK-33 treated infected samples (Treated virus) (green).

**Figure S8.**
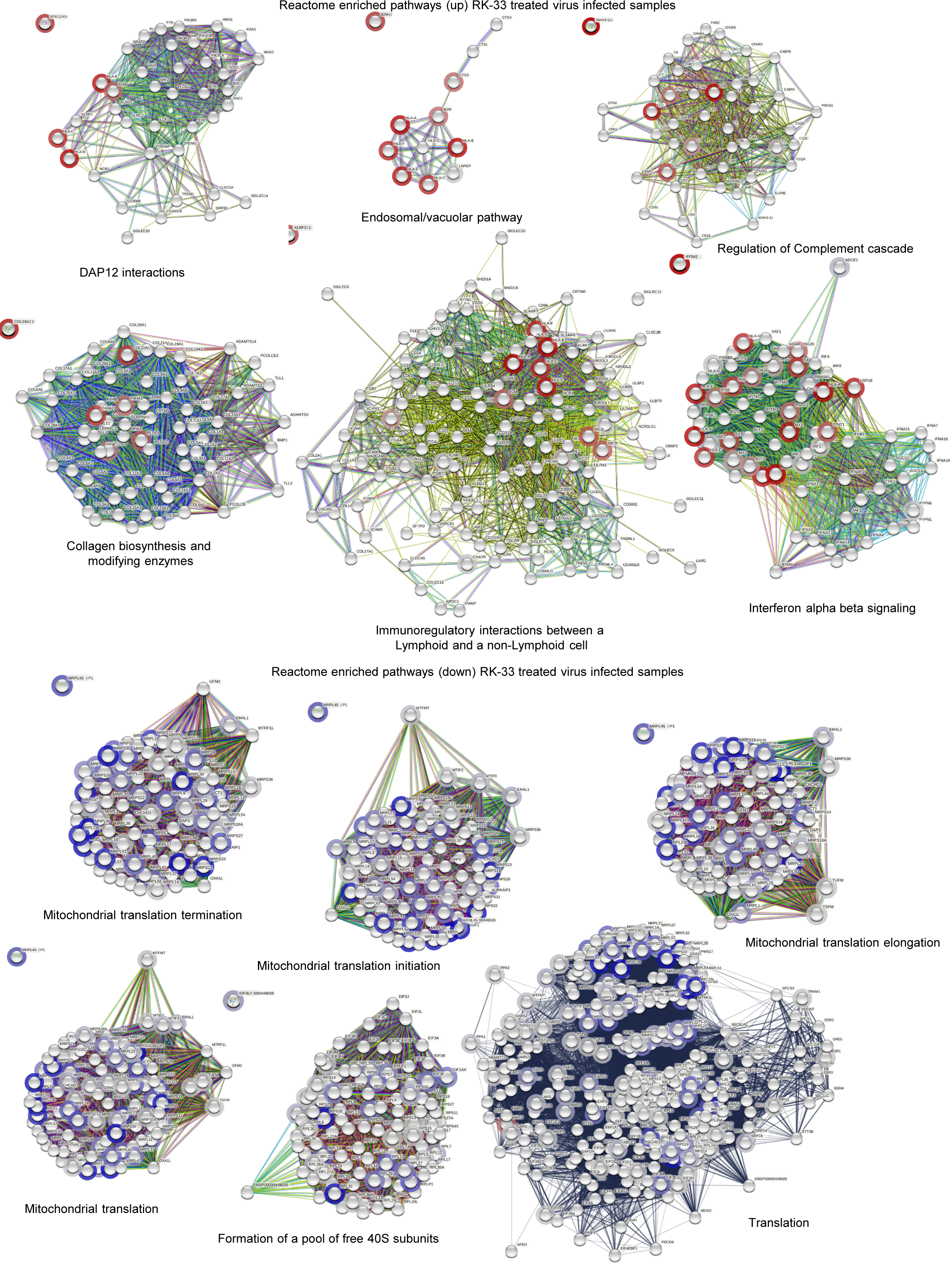
STRING analysis of proteins showing top six Reactome enriched (up and down) pathways of RK-33 treated virus infected samples.

**Figure S9.**
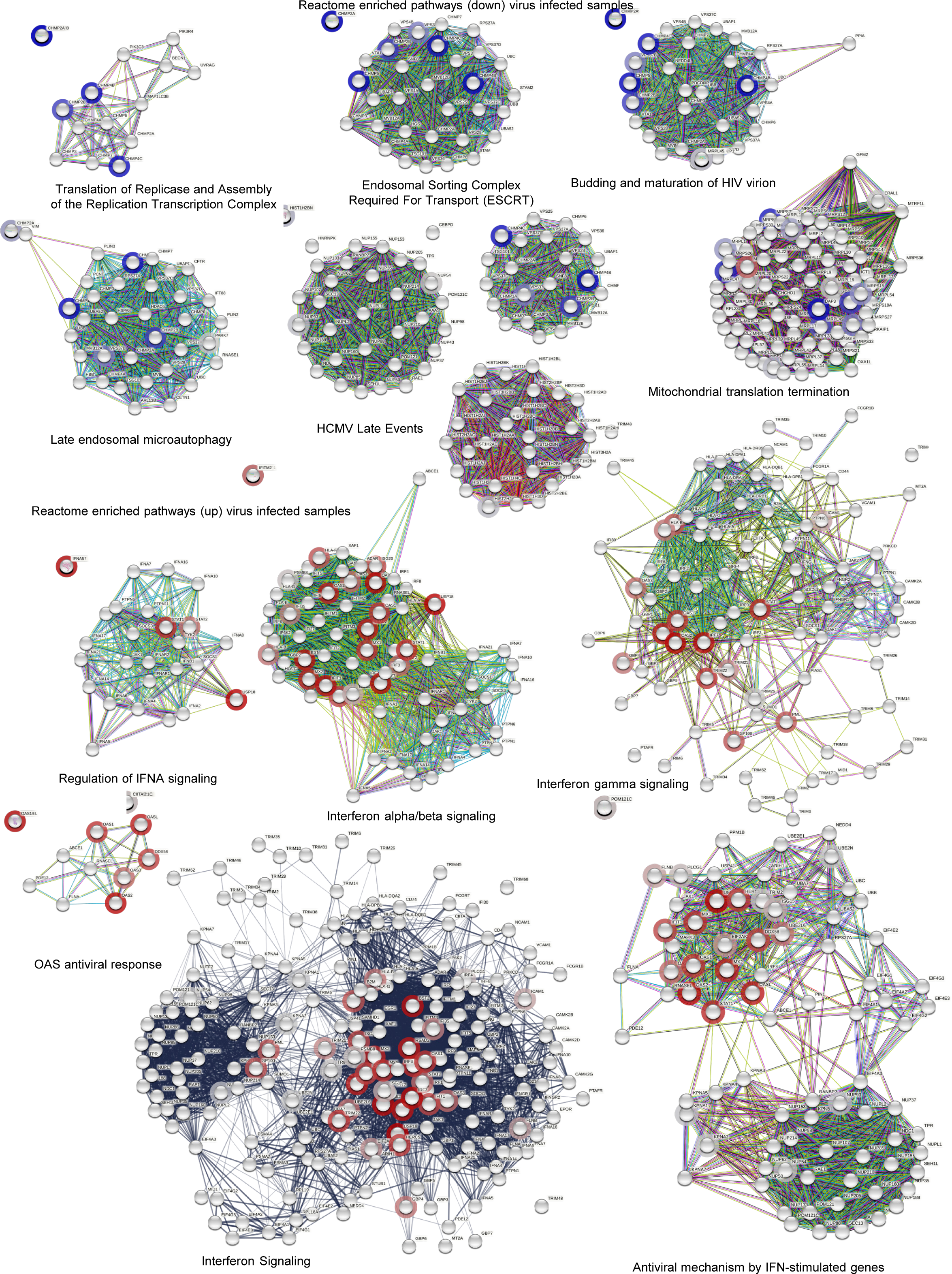
STRING analysis of proteins showing top six Reactome enriched (up and down) pathways of virus infected samples.

**Table S1.**
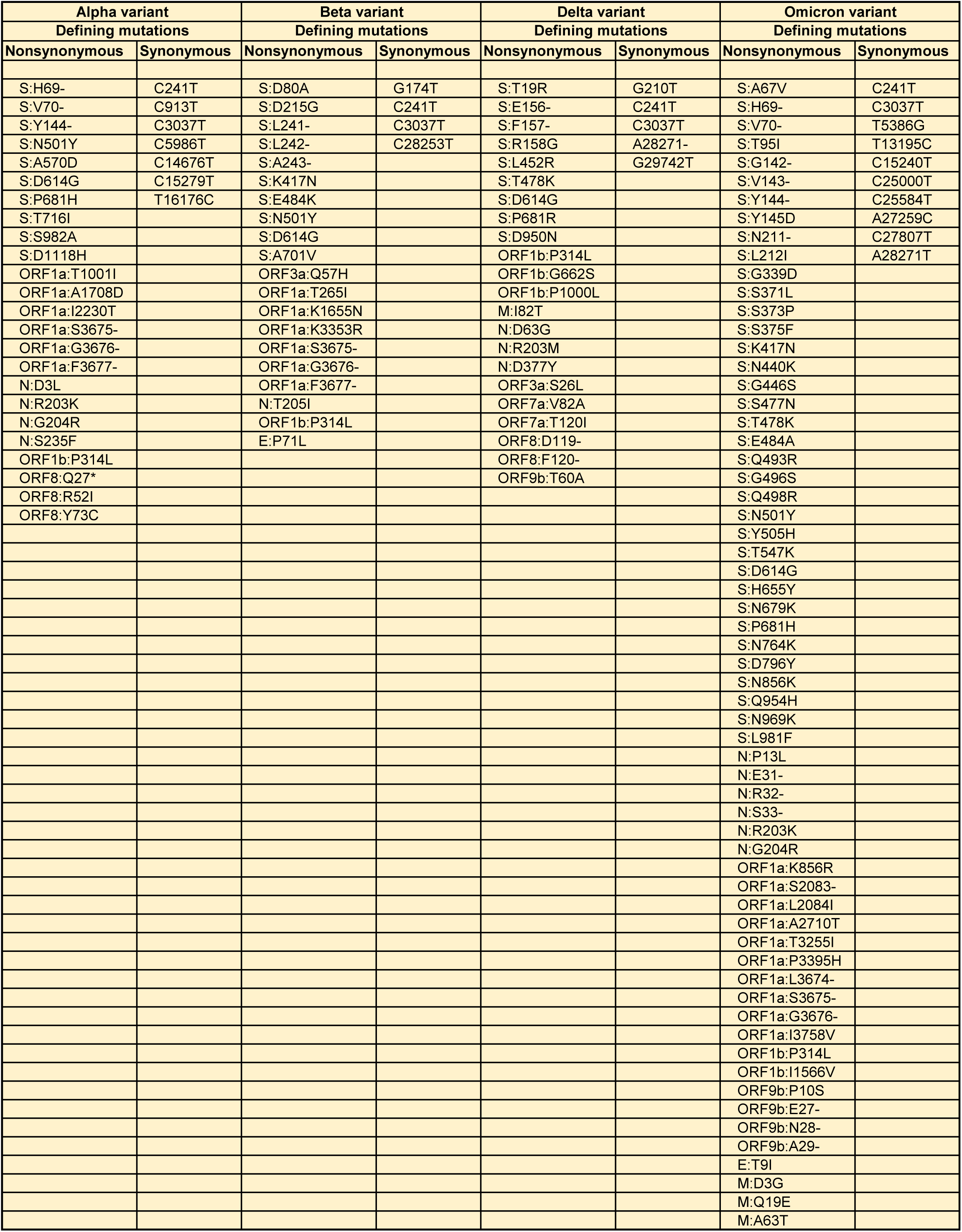
A list of defining mutations present in the Alpha, Beta, Delta, and Omicron variants of SARS-CoV-2.

## Notes

### Competing Interest Statement

Venu Raman holds a patent on the composition of RK-33 (US patent # 8,518,901).

